# Horus: Reproducible Workflow Manager for Biomolecular Modelling

**DOI:** 10.1101/2025.09.26.678751

**Authors:** C Domínguez-Dalmases, A Cañellas-Solé, R Lambea-Jané, M Municoy, V Guallar

## Abstract

Horus is a versatile, multi-platform reproducible-workflow management system designed for biomolecular modelling and computational life sciences. Available as both a desktop application and a web-based service, Horus offers researchers an intuitive graphical interface alongside a robust Python API, democratising the creation, execution, and management of complex scientific workflows. Its modular architecture uses individual blocks that can be easily customised and extended, allowing for an integration with diverse computational tools and resources. Horus supports remote execution via SSH to efficiently dispatch tasks to high-performance computing clusters in remote machines. The platform also features integrated 3D and 2D molecular visualizers for interactive structure analysis, which enhances the interpretability of simulation results. Its design emphasises reproducibility, scalability, and user-friendliness, positioning Horus as a valuable tool to advance research in structural bioinformatics and related fields.

## Introduction

The rapid expansion of computational tools in biomolecular modelling has introduced significant challenges to researchers. Managing all the functionalities, frequent updates, and diverse dependencies of these tools often requires substantial technical expertise. In addition, integrating disparate applications, each potentially operating in unique environments or on separate machines, into cohesive workflows remains a complex task.

Existing workflow management systems offer partial solutions. For example, Galaxy (1, 2) provides a user-friendly graphical interface, facilitating access to a wide range of bioinformatics tools. However, its installation can be cumbersome for users without advanced IT skills, and its web-based nature may not suit all computational environments (e.g., those with limited internet access or strict security protocols). Conversely, Nextflow (3) offers greater flexibility and scalability, supporting execution across various platforms, including local machines, clusters, and cloud services. Nevertheless, it lacks a graphical user interface, necessitating familiarity with its domain-specific language, which can be a barrier for some users, even when using cloud-based solutions like the Seqera platform^1^ or nf-core Launch^2^, which offer limited visualization capabilities.

To address these limitations, we introduce Horus, a free versatile reproducible-workflow management system designed for biomolecular modelling tasks, but not limited to them. Horus combines an intuitive graphical user interface with a powerful Python (4) API, enabling users to create, manage, and execute complex workflows effortlessly. Its modular architecture supports the integration of additional tools, regardless of their underlying environments or locations. Furthermore, Horus offers both standalone and web-based deployments, providing flexibility to accommodate varying requirements and collaborative needs. By bridging the gap between usability and functionality, Horus aims to enhance the efficiency and reproducibility of computational research in the life sciences.

### Application Overview

Horus is built using standard web technologies, ensuring compatibility across various operating systems and devices. This design choice facilitates both local and remote deployments, allowing users to operate Horus as a standalone desktop application or access it via a web browser through a centralised server.

#### Modular

At its core, Horus employs a modular framework that supports the integration of diverse computational tools and resources. The system’s extensibility is achieved through the HorusAPI, a robust Python API that allows users to incorporate new functionalities. This modularity ensures that Horus can adapt to the evolving needs of the scientific community, accommodating a wide range of bioinformatics applications.

#### Remote

To enhance computational efficiency, Horus offers native support for secure remote execution via SSH, allowing tasks to be dispatched to high-performance computing clusters.

Integration with job scheduling systems like SLURM (5) further optimises resource management, ensuring that computational workloads are handled effectively.

#### Visual

Visualisation is a key component of Horus’s functionality. The platform integrates open-source tools such as Mol* (6), OpenBabel (7), and JSME (8), providing users with interactive molecular structure visualisation and editing capabilities. These integrations enable real-time analysis and interpretation of complex biomolecular data, enhancing the overall user experience.

#### Intuitive

By combining a user-friendly interface with powerful computational capabilities, Horus is positioned as a comprehensive solution for researchers to manage and execute complex bioinformatics workflows efficiently. Moreover, Horus aims to democratise the development of graphical user interfaces (GUIs), making it easier for researchers to incorporate intuitive front-ends into their own software tools. For example, the developers of PandaDock^3,4^ used Horus to create a functional GUI without requiring extensive additional development.

### Workflow Management and Visualisation Tools

Horus offers an intuitive and interactive user interface designed to facilitate the construction and execution of complex biomolecular workflows. This interface consists of an infinite 2D canvas with a drag-and-drop functionality for the assembly of blocks into the workflow. Users can establish connections between these blocks, defining the sequence and logic of computational tasks. Upon execution, Horus automatically manages input dependencies and orchestrates the execution sequence, ensuring efficient workflow progression.

For molecular visualisation, Horus integrates the Mol* engine: a high-performance WebGL-based (9) tool known for its application in the RCSB Protein Data Bank (10). Mol* supports the rendering of large macromolecular structures, including proteins, nucleic acids, and complexes, offering features such as coarse-grained visualisation and interactive selection of molecular components. Within Horus, Mol* is tightly connected to the workflow environment, enabling blocks to access and manipulate molecular data directly, as seen in Figure 1. Users can define specific regions of interest, such as atoms, residues, chains, or spatial selections like spheres and boxes, through intuitive interactions within the visualizer.

**Fig. 1:**
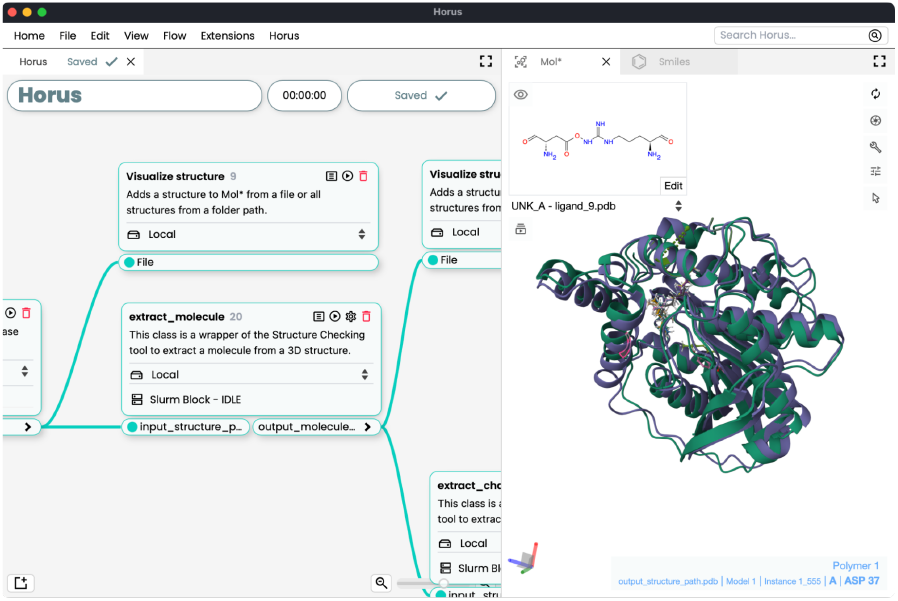
Example of a Horus workflow with the Mol* visualisation panel open. On the left, individual workflow blocks are shown with their inputs and outputs connected, illustrating the data flow between computational steps. On the right, the Mol* panel displays molecular structures generated as outputs of the workflow, enabling real-time visualisation and interactive analysis of the results.

Complementing the 3D visualisation, Horus incorporates a 2D molecular editor powered by JSME. This editor allows users to import molecular data in formats like CSV, SMI, and SDF, converting them into editable SMILES (11) representations. JSME facilitates the drawing and modification of chemical structures, with real-time updates reflected in an accompanying data table. Users can annotate molecules with custom properties, sort and group entries, and visualise associated data through interactive plots. Additionally, the system supports the interconversion between SMILES and SDF formats, enabling seamless transitions between 2D editing and 3D visualisation.

Horus’s modular architecture permits the integration of custom visualisation components. Developers with proficiency in web technologies can embed HTML, CSS, and JavaScript-based views into the platform through the Horus API Extensions() class.

This flexibility allows the improvement of data interpretation and presentation. Apart from that, Horus comes with several default visualization views for diverse data types, including CSV files, PDFs, images, text, and HTML content.

Workflows are encapsulated in.flow files, which store block configurations, the state of the 3D molecular visualizer, and any molecules present in the 2D SMILES editor. These files are easily shareable, allowing peers to review results, replicate analyses, or modify workflows for extended studies. This allows Horus workflows to be highly reproducible, as if it worked in one environment, the workflow must work into another given the same conditions. In Figure 2, a diagram of how the Horus workflow engine works is shown.

**Fig. 2:**
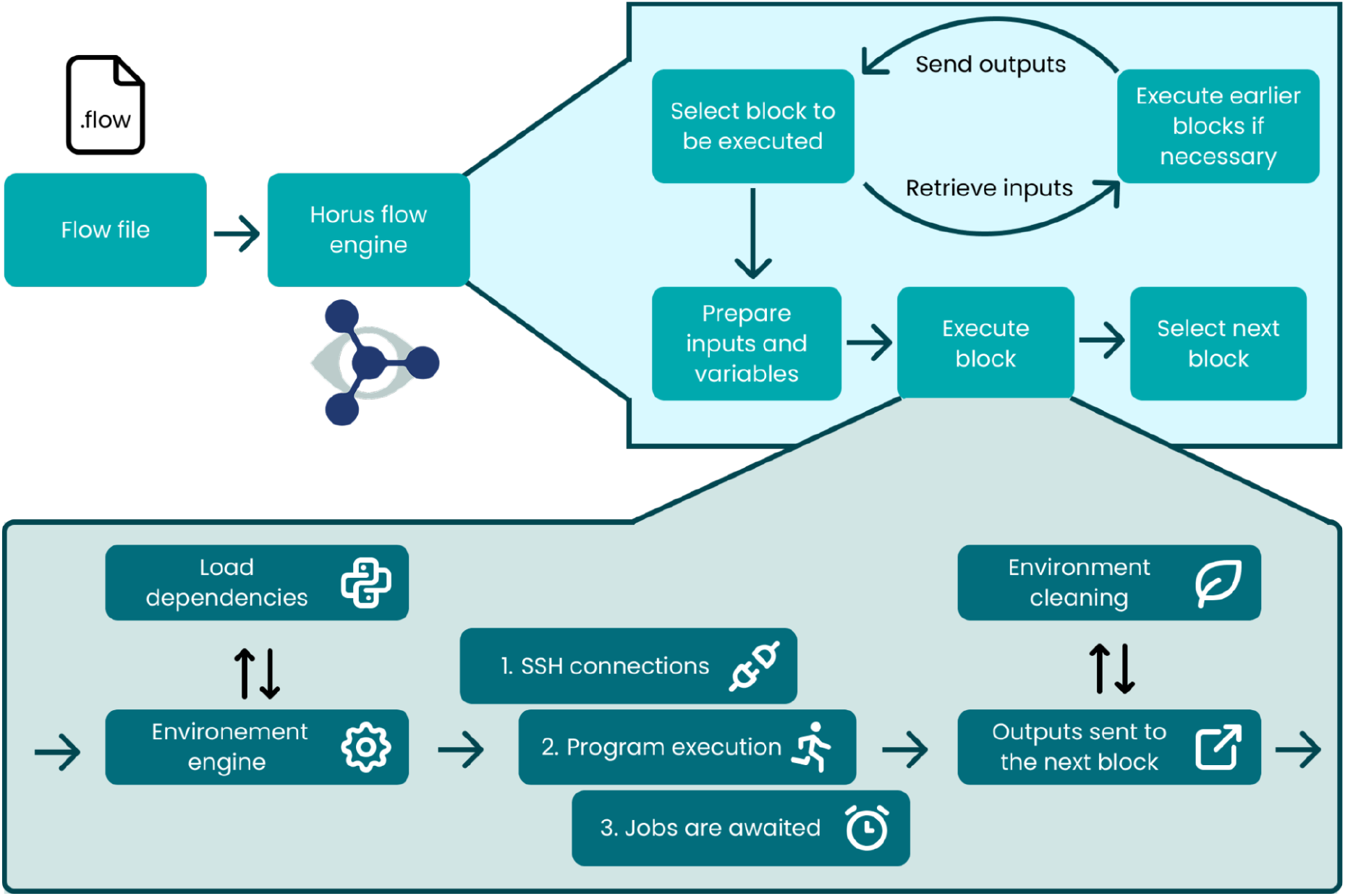
The Horus workflow engine starts by reading the workflow file and instantiating the blocks placed by the user. Next, the engine finds the block designated by the user to be executed and identifies any inputs that the block may need. If any input that comes from another block is missing, the selected block to be executed will change to that input block, and the process will start from the beginning. This works iteratively until all dependencies needed to run the initially selected block are fulfilled. During the preparation of the inputs phase, the outputs of previous blocks are assigned to the inputs of the selected block, ensuring that the execution of the block contains all the expected values. Before the block starts running, the environment engine pre-loads a small Python environment in memory using an in-house implementation of the Python module system. Once the environment is ready, the block connects to any remote (if selected) and runs the code of the block and any associated programs. Finally, if no SLURM jobs are pending, the environment is cleaned and the outputs are sent to the next block.

### Remote Execution and Resource Management

To support computational workloads that span across local and remote environments, Horus integrates secure remote execution capabilities via the widely used Python library Paramiko^5^. This enables each workflow block to independently establish an SSH (12) connection to a configured remote machine using standard credentials (username, host, and either password or private key). Once connected, blocks can upload and retrieve files, execute commands, and interact with remote job scheduling systems—all from within the Horus interface. An example of a workflow that includes 3 different runtime environments (local, and two separated remote servers) is explained in Figure 3.

**Fig. 3:**
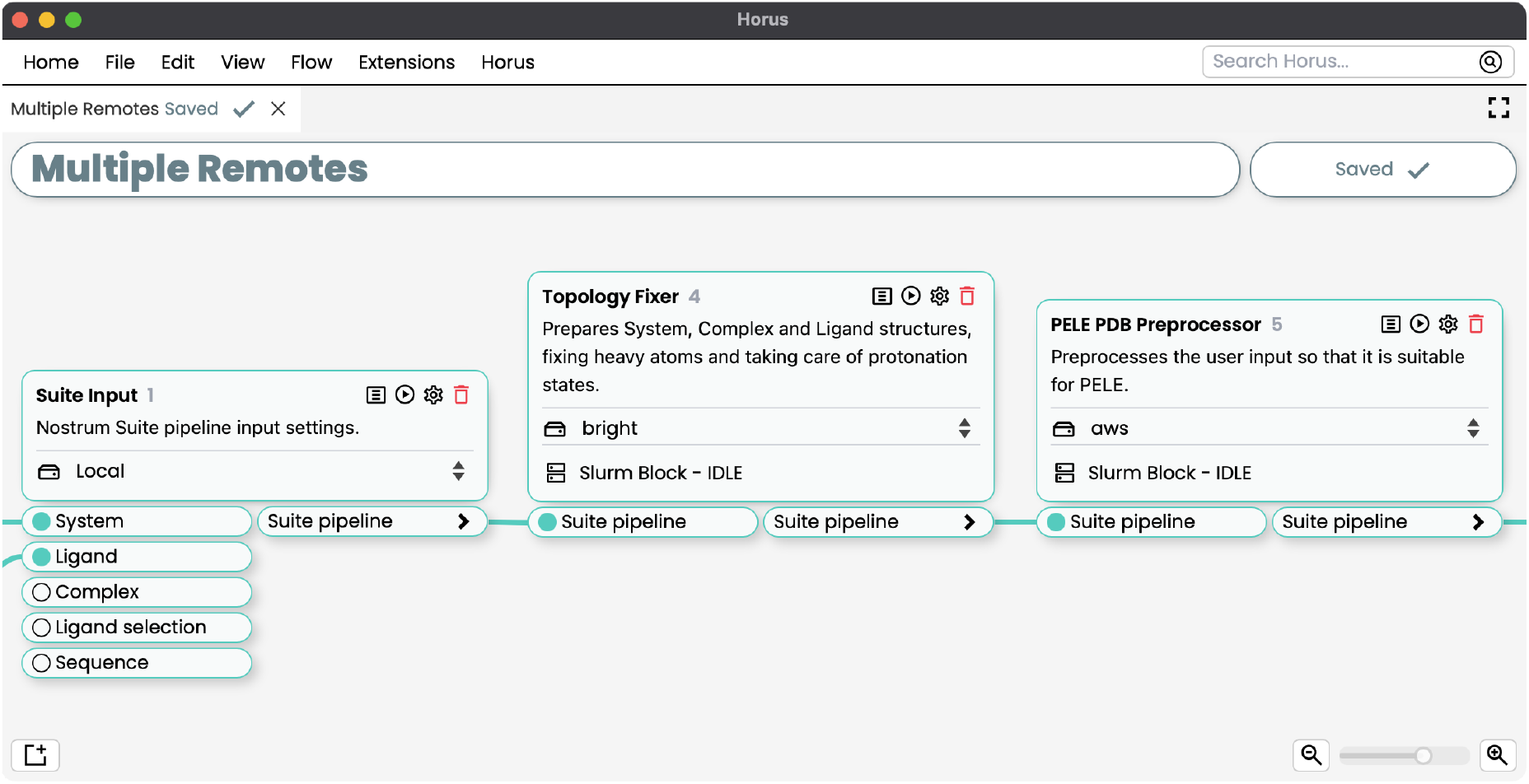
In this workflow diagram, using the Nostrum Suite plugin, three consecutive blocks are executed across different remotes. The first block, **Suite Input**, runs on the local machine, where commands and SLURM jobs are executed. The outputs of this block are then passed to the second block, **Topology Fixer**, which connects via SSH to the remote named **bright**. After establishing the connection, the selected input files are uploaded, commands and SLURM jobs are executed remotely, and after completion, the relevant results are downloaded back to the local workflow directory. Finally, the results are forwarded to the third block, **PELE PDB Preprocessor**, which repeats the same process on the remote named **aws**, ensuring that inputs are uploaded, jobs executed remotely, and outputs returned locally. This setup allows the workflow work across multiple remote machines while keeping the local environment synchronized with all intermediate results.

The ability to run simulations across configured remotes is one of the key features that makes Horus stand out from other workflow managers or runners. This way, a workflow can be easily automated across environments which can be easily run by a simple few clicks.

High-performance computing (HPC) integration in Horus is centred around support for SLURM, a widely adopted open-source job scheduler. Users can configure blocks to submit one or multiple SLURM jobs concurrently, with each job tracked through its assigned job ID on the target cluster. Horus provides a built-in monitoring interface where users can inspect the submission script, view real-time stdout/stderr logs, and track job state changes as they progress through the queue.

Furthermore, Horus fully supports SLURM job status feedback: execution of the next blocks is stopped until all associated jobs complete. If a job fails, the system prompts the user whether to resubmit the job or continue even if some jobs failed. This approach provides transparency in managing long-running tasks.

### API and Extensibility

Horus is designed to be highly modular with a Python and web-based plugin system that allows users to expand the platform to their needs. In Horus, plugins are the main way to extend functionality. They can provide blocks, custom interfaces, and predefined workflows. This extensibility system is called the HorusAPI^6^.

Plugins are loaded at startup and support two creation methods: automatic via the *create-horus-plugin* command or by structuring a plugin folder with a *plugin*.*meta* JSON file describing its metadata. Plugins can include PIP dependencies, private or pre-installed libraries inside an *Include* folder, inter-plugin imports and install/uninstall scripts (preinst.sh, postinst.sh, prerm.sh, postrm.sh) for custom setup behaviour.

Developers can enable *Development Mode* in the app to reload plugins without restarting. *Symlinks* can be used to point the Horus plugins folder to a development directory for rapid iteration. In the HorusAPI, Blocks constitute the fundamental execution units, each responsible for carrying out a predefined *Action* based on a set of associated **PluginVariable** instances, as seen in Figure 4. These blocks serve as modular wrappers around Python functions and are classified into three types depending on their behaviour: **PluginBlock**, the most common type, executes local functions; **InputBlock** is used for data injection via a single input variable; and **SlurmBlock**, which is designed for distributed computation, encapsulates both a pre-submission and post-completion function for SLURM-based job execution. While **PluginVariable** objects model input/output parameters, the dynamic values should be retrieved within the *Action* function through the *block*.*variables, block*.*inputs*, or *block*.*config* mappings, using the variable identifiers as keys. Outputs are defined programmatically via *block*.*setOutput()*. Configuration parameters can be declared through the **PluginConfig**, a subclass of **PluginBlock**, and are consistently accessible and mutable across plugin executions. Moreover, developers can persist arbitrary runtime metadata using the *extraData* property, available at both the block and flow levels. This enables context-aware logic across repeated executions, though its lifecycle must be explicitly managed. Errors encountered during execution beyond the *Action* function’s scope will automatically mark the block as failed, triggering a user-facing alert in the graphical interface.

**Fig. 4:**
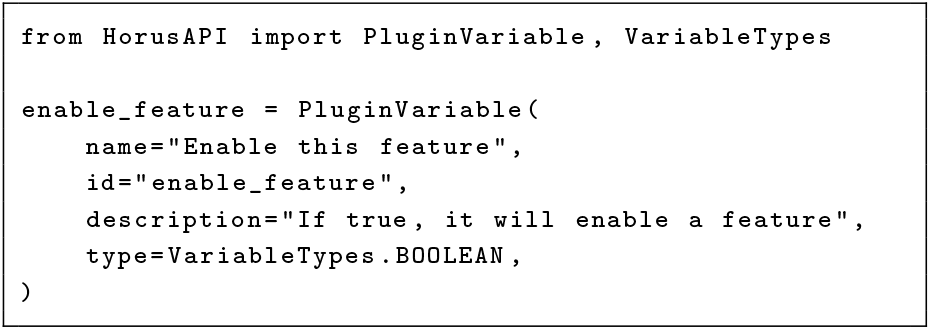
Example of creating a Boolean plugin variable using the HorusAPI.

In summary, the HorusAPI offers a simple and accessible way to build blocks and extend functionality. For developers who are familiar with Python, HTML, CSS, and JavaScript, Horus allows embedding custom views, developing custom blocks, and implementing unique integrations. Developers may install the HorusAPI directly from PyPI (with **pip install HorusAPI**). Nevertheless, the HorusAPI allows non-technical users to develop their custom plugins, providing a clear and simple Python interface to the graphical user interface components.

### Plugin Repositories

Plugins developed for Horus can be published and shared through the public Horus Plugin Repository, accessible at https://horus.bsc.es/repo and from the **Horus Plugin Manager** inside the application, as seen in Figure 5. This platform facilitates the distribution of community-driven plugins, enabling researchers to extend Horus with minimal effort through a graphical interface.

**Fig. 5:**
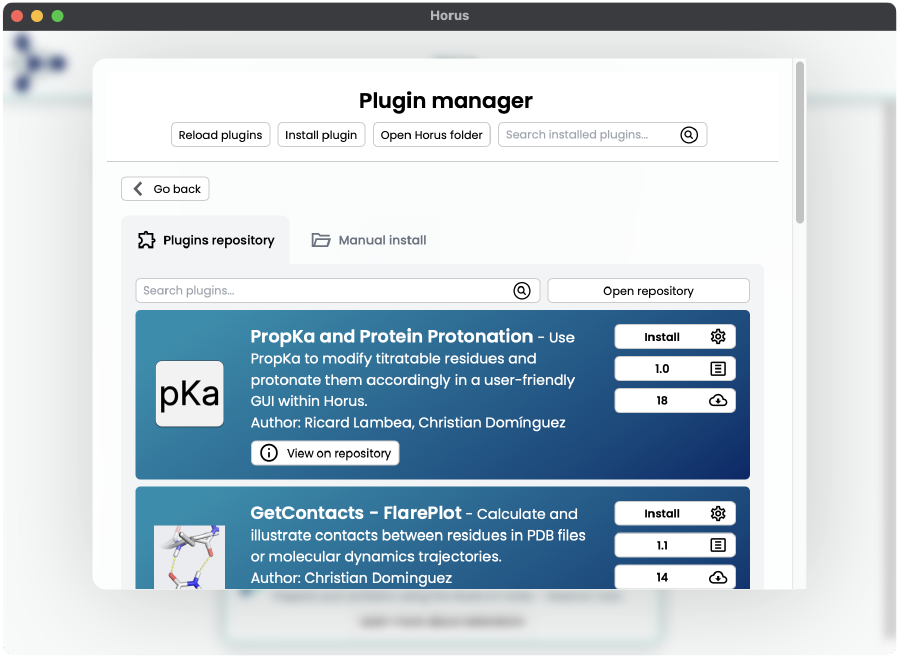
The Horus Plugin Repository interface, integrated directly inside the application. Users can browse, search, and install plugins with just a few clicks. Each plugin entry displays the number of downloads, the current version, and a direct link to the corresponding repository page, where additional documentation and usage information are available.

Notable plugins currently available include BioExcel BioBB Blocks (13), which include open source tools like Gromacs (14), OpenBabel (7), PyTorch (15) and commercial software like Amber (16, 17). Moreover, standalone rDock (18), PandaDock, ESMFold (19), ProteinMPNN (20), PropKa (21, 22), Nostrum Suite^7^ (which includes PELE (23)), PyDock (24) and the GetContacts^8^ library plugins are available. Developers may upload their packaged plugins in.**hp** format directly to the repository or distribute them manually. The latest version of Horus supports custom repositories. Enabling groups and companies to publish their packages on their own infrastructure.

### Use Cases and Applications

Horus has already demonstrated its utility across a diverse set of scientific software environments, enhancing both accessibility and reproducibility. Several key applications—including our EAPM (Electronic and Atomic Protein Modelling) lab tools, Nostrum Suite, GeoDirDock, BioExcel BioBB workflows, and various lightweight utilities—have been successfully integrated into Horus through dedicated plugins, as mentioned before.

In the case of the EAPM tools, Horus enabled the full integration of a protein engineering pipeline, facilitating streamlined execution across preparation, computation, and result visualisation stages. This workflow, which previously required manual coordination across tools and scripts, is now fully automated within a reproducible and shareable graphical environment.

Nostrum Suite uses Horus to encapsulate and distribute all of its computational tools via an intuitive graphical user interface. This significantly lowered the barrier for adoption among collaborators and clients and allowed for rapid delivery and visualisation of computational results.

The BioExcel Building Blocks (BioBB) have also been fully ported to Horus, with each block supporting distinct execution modes: singularity, docker or conda, local or remote and SLURM or subprocess. These blocks are readily accessible through the Horus Plugin Repository.

#### Case Study: Protein Engineering of 5JD4 Enzyme using Horus

A representative use case that demonstrates the capabilities of Horus is the protein engineering pipeline available on the official Horus website^9^. Protein engineering is a vital strategy for optimising protein function or introducing novel properties. Horus facilitates the construction of advanced, modular workflows that integrate multiple tools across the protein design and analysis pipeline. In this specific example, we target the enzyme 5JD4 (25). This enzyme represents the Ser161Ala mutant of the LAE6 enzyme—a mutation that underscores the catalytic role of the Ser161 residue (26).

The workflow incorporates ProteinMPNN for sequence design, ESMFold for structure prediction, and rDock for molecular docking. Prior to these core steps, preprocessing of the 5JD4 structure is performed using BioExcel’s BioBB blocks and Biopython (27). Horus’s flexibility allows integrating heterogeneous tools into a unified visual pipeline, as seen in Figure 6. Moreover, a tutorial in PDF format is embedded within the application, open-able as an additional panel thus improving cohesion inside the platform.

**Fig. 6:**
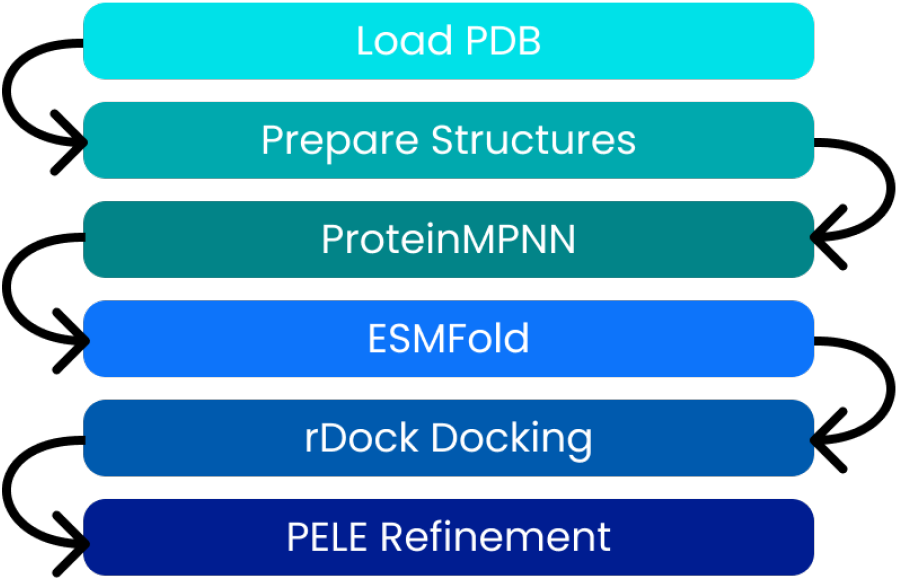
Enzyme engineering workflow diagram. ProteinMPNN, ESMFold and rDock operate alongside each other, enabling an integrated pipeline for sequence design, structural prediction, and molecular docking. This cohesive interface allows researchers to combine multiple bioinformatics tools within a unified graphical environment, streamlining the entire protein engineering process. The detailed workflow interface inside Horus can be previewed at https://horus.bsc.es/pep. PELE blocks are not shown in the public Horus instance as they are not freely available and require a license.

The study focuses on introducing rational and automated mutations into the pocket site of 5JD4 to evaluate their impact on enzymatic activity. Two specific substitutions, derived from the literature, are introduced: Glu27→Asp (E27D) and Leu216→His (L216H). ProteinMPNN is further used to propose additional mutations in the catalytic region, targeting stability or enhanced function.

To assess the functional consequences of these mutations, the ligand ethyl benzoate (CCC(=O)OC1=CC=CC=C1) is docked into both the wild-type and engineered protein structures using rDock. The docking strategy first establishes a baseline score with the native 5JD4, followed by comparative evaluation against the mutated variants. Differences in docking scores provide insights into the influence of each mutation on substrate binding affinity and potential enzymatic performance. After performing the docking simulations, we can take the outputs and visualise them inside of Horus, using the “Visualize structure block”. We can simultaneously compare the mutated and the wild-type results inside the same workflow with just a few clicks. Finally, we could perform some induced fit docking with PELE, in order to verify the results. Using the Nostrum Suite Pro plugin for Horus, allows us to quickly set up a PELE simulation in seconds. We can just open one of the provided templates, and set as the inputs the obtained results from rDock. Moreover, the Nostrum Suite Pro plugin includes an interactive results visualisation interface, designed specifically for Horus. Using the multi-pane visualisation capabilities of the application, one can compare both simulations, the mutated and the wild type, at the same time, as seen in Figure 7.

**Fig. 7:**
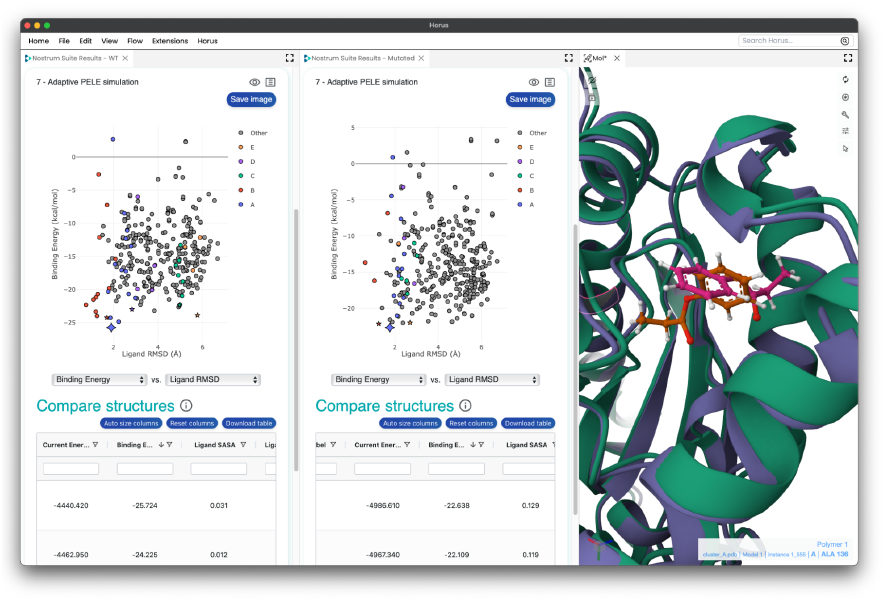
Multi-pane view of the Nostrum Suite Pro results. On the left is the energetic profile obtained from a PELE simulation of the wild type. In the centre is the energetic profile of the mutated protein. Finally, the Mol* viewer with the best poses for each structure is displayed. Clicking on a point in the plot will reveal in the viewer the corresponding 3D structure of the specified pose.

This case study exemplifies how Horus enables reproducible, extensible, and GUI-accessible scientific workflows by seamlessly orchestrating diverse computational tools within a single plugin ecosystem.

#### Case study: Predicting potential 1UDI binding sites with pyDock ODA

Another great use case of software integration as an Horus plugin is the pyDock ODA (Optimal Desolvation Area) plugin, developed at Nostrum Biodiscovery.

PyDock ODA is a specific module from the pyDock software which analyzes optimal desolvation patches on a protein surface to predict potential interfacial binding sites.

In the current example, the input used is the protein 1UDI (28), in PDB format. In Figure 8, it can be seen the protein acting as ligand (light blue cartoon), and the receptor protein (yellow cartoon), onto which its ODA predicted result has been superposed as a colored molecular surface representation.

**Fig. 8:**
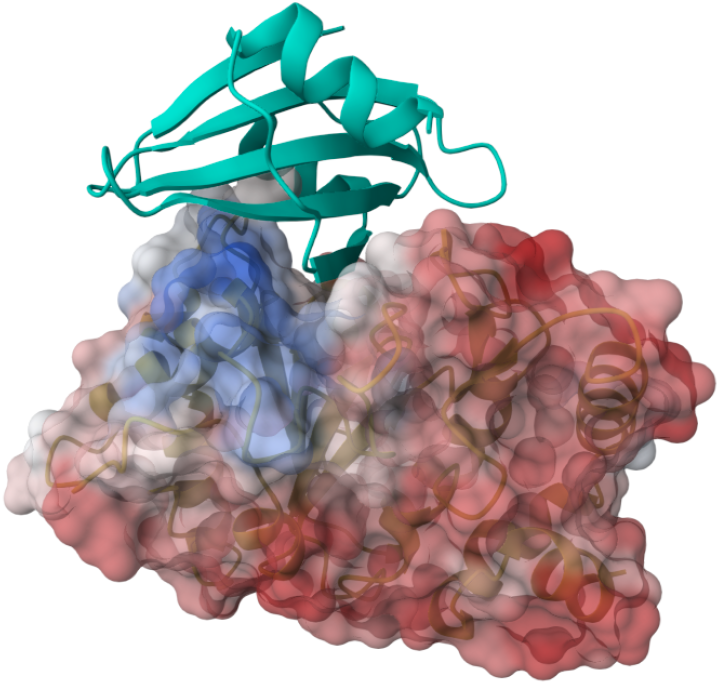
pyDock ODA structure representation. Using the HorusAPI, the pyDock ODA block can load and display the structure the Mol* viewer with a custom visual schema programmatically.

As stated by the pyDock authors in the pyDock ODA documentation^10^: “Regions predicted to belong to an interface have an ODA value below −10”. Thus, the colour scale of the molecular surface representation takes values from −15 to 5 in a blue-white-red palette. This can be interpreted in the following way: blue coloured residues have an ODA predicted value shifted to the negative side of the scale, residues which have a value between the two value range limits are displayed in white, and eventually, residues showing a more positive predicted value are coloured in red, meaning areas predicted as unlikely to be potential binding interface sites.

This is achieved using the Horus **MolstarAPI**. In Figure 9, an example of such implementation is depicted. The **addMolecule** method takes a **MolstarThemeOptions** object, which is passed to the theme argument of the method. At the same time, this object definition takes several keyword arguments: representation, which defines the structure design in Molstar (e.g. molecular surface); colour, which allows to define the colour theme for the current structure, i.e. the property (**uncertainty** column from ODA resulting PDB file in this case) which will be used to color the structure; and **colorParams**, which allows to customize how the color theme is displayed onto the representation. In this case, by specifying a range of values ([-15, 5]) to the domain key, we can display a palette of three colours, which correspond to the predicted ODA values.

**Fig. 9:**
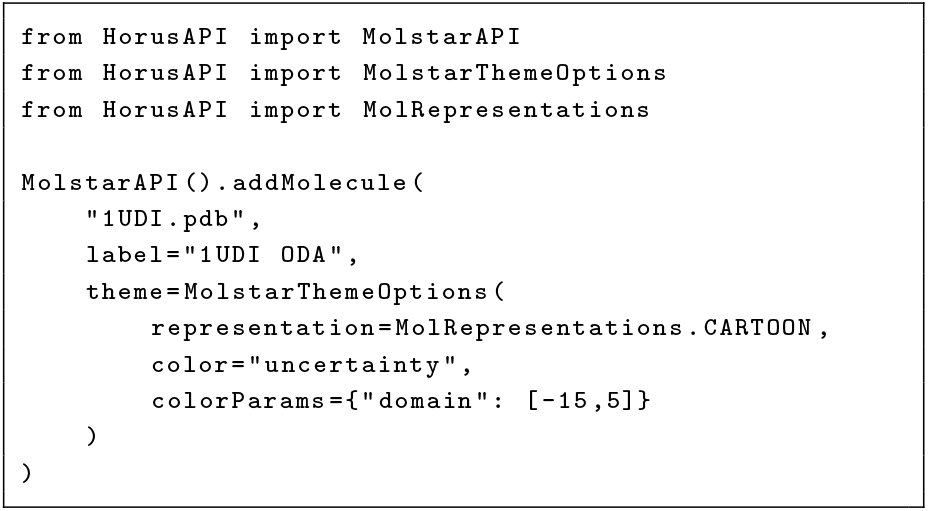
Example usage of the MolstarAPI within the pyDock plugin ODA block, showing how to load a PDB structure (1UDI.pdb) and visualize it with a cartoon representation coloured by uncertainty (ODA values).

## Data Availability

Horus can be freely downloaded from https://horus.bcs.es. A demo is available free to use at https://horus.bsc.es/demo. The protein engineering pipeline demo can be found at https://horus.bsc.es/pep.

Horus is designed following the FAIR (29) principles to ensure that data, workflows, and plugins are Findable, Accessible, Interoperable, and Reusable. Resources within Horus are assigned unique identifiers and complemented with rich metadata to facilitate discovery. The platform provides open access using standard protocols and maintains metadata availability independent of data status. Horus uses standardised, machine-readable formats and common vocabularies to enable integration with other tools. Additionally, clear licensing, and documentation, following community standards, support reproducibility and reuse of workflows.

---

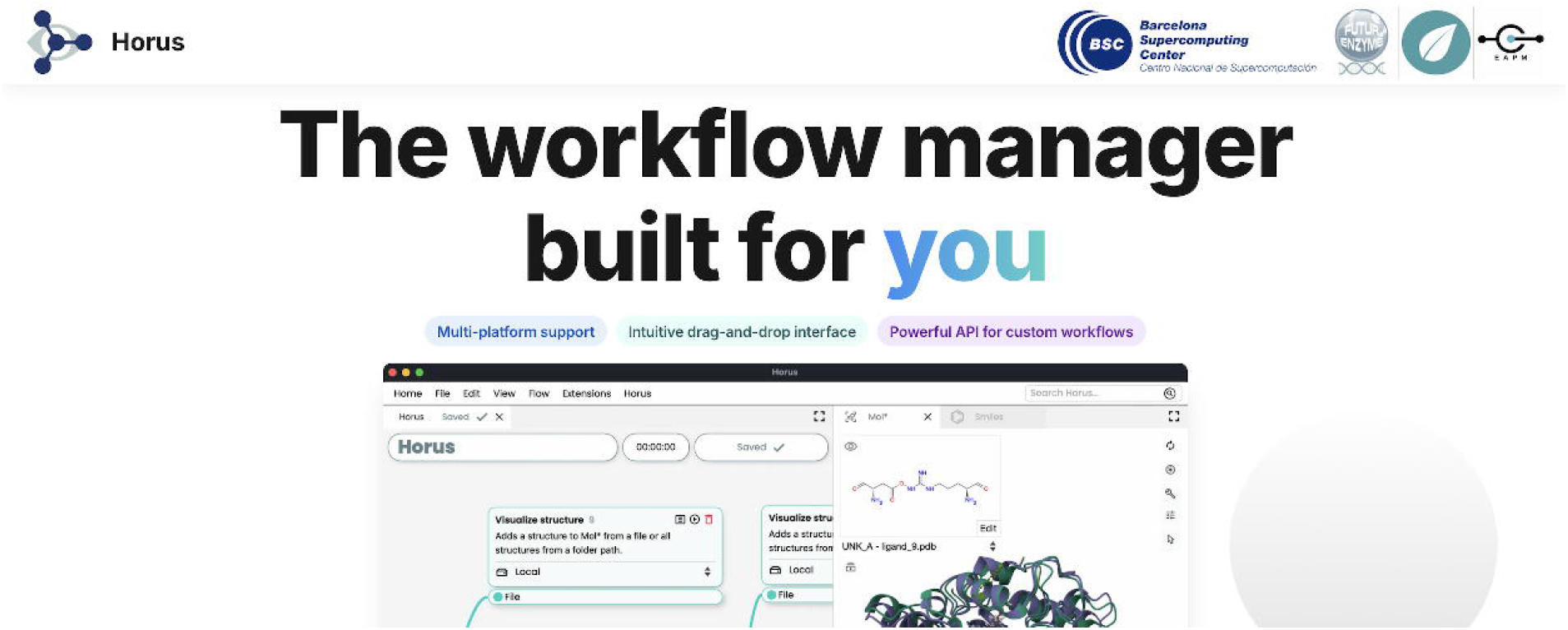

## Footnotes

1 Seqera: https://seqera.io

2 nf-core Launch: https://oldsite.nf-co.re/launch

3 PandaDock: github.com/pritampanda15/pandadock

4 pandadock horus: github.com/chdominguez/pandadock_horus

5 Paramiko: https://github.com/paramiko/paramiko

6 The HorusAPI documentation is available at https://horus.bsc.es/docs/developer_guide/

7 *Nostrum Suite*. Nostrum Biodiscovery S.L., Barcelona, Spain, 2025. Available at: https://docs.nostrumbiodiscovery.com

8 *GetContacts*. Available at: https://getcontacts.github.io/

9 Protein Engineering Pipeline: https://horus.bsc.es/pep

10 https://life.bsc.es/pid/pydock/doc/tutorial.html#introduction

